# Soft sweeps predominate recent positive selection in bonobos (*Pan paniscus*) and chimpanzees (*Pan troglodytes*)

**DOI:** 10.1101/2020.12.14.422788

**Authors:** Colin M. Brand, Frances J. White, Nelson Ting, Timothy H. Webster

**Author notes:** Corresponding Author: Colin M. Brand, University of Oregon, Department of Anthropology, 1218 University of Oregon, Eugene, OR 97403.

## Abstract

Two modes of positive selection have been recognized: 1) hard sweeps that result in the rapid fixation of a beneficial allele typically from a *de novo* mutation and 2) soft sweeps that are characterized by intermediate frequencies of at least two haplotypes that stem from standing genetic variation or recurrent *de novo* mutations. While many populations exhibit both hard and soft sweeps throughout the genome, there is increasing evidence that soft sweeps, rather than hard sweeps, are the predominant mode of adaptation in many species, including humans. Here, we use a supervised machine learning approach to assess the extent of completed hard and soft sweeps in the closest living relatives of humans: bonobos and chimpanzees (genus *Pan*). We trained convolutional neural network classifiers using simulated data and applied these classifiers to population genomic data for 71 individuals representing all five extant *Pan* lineages, of which we successfully analyzed 60 individuals from four lineages. We found that recent adaptation in *Pan* is largely the result of soft sweeps, ranging from 73.1 to 97.7% of all identified sweeps. While few hard sweeps were shared among lineages, we found that between 19 and 267 soft sweep windows were shared by at least two lineages. We also identify novel candidate genes subject to recent positive selection. This study emphasizes the importance of shifts in the physical and social environment, rather than novel mutation, in shaping recent adaptations in bonobos and chimpanzees.

## Introduction

The identification of adaptative traits and their genetic basis is one of the central goals of evolutionary biology. Two approaches, top-down and bottom-up, have been used to accomplish this goal; the latter of which leverages population-level data to recognize the genomic signatures of positive selection (Barrett and Hoekstra 2011). At the genomic level, the process of adaptation results in a window of reduced variation that erodes over time. As these signatures do not persist, they can only be used to infer selection over a particular time scale in a population. In most species, this time frame is restricted to a few thousand generations, roughly ~ 200,000 years in humans (Oleksyk et al. 2010). The classic model for positive selection for a given locus proposes that a single, novel mutation, that confers a fitness advantage (i.e., a beneficial allele) will rapidly spread in a population and eventually reach fixation (Maynard Smith and Haigh 1974). Neutral polymorphism adjacent to the novel allele will ‘hitchhike’, resulting in a distinct pattern of reduced genomic diversity at the locus and surrounding sites. The term ‘hard sweep’ has been used to identify this pattern and process.

‘Soft sweeps’ describe the presence of two or more haplotypes that occur at intermediate frequencies (Hermisson and Pennings 2005). Thus, the signature of a soft sweep is intermediate to those of neutral or ‘background’ genomic variation and the signature of a hard sweep. This pattern can result from recurrent *de novo* mutations following positive selection. Alternatively, soft sweeps can also result from positive selection on standing genetic variation where alleles were already present in a population before selection. This variation may be the result of independent mutations (multiple origin soft sweep) or when an adaptive allele arose before selection, but multiple copies have subsequently swept through the population (single origin soft sweep). Soft sweeps are often incorrectly viewed synonymously with standing genetic variation; hard sweeps can emerge from standing genetic variation if a single copy of the beneficial allele was the ancestor of all beneficial alleles in a sample (Hermisson and Pennings 2017).

Hard and soft sweeps are locus-specific and, thus, not mutually exclusive across a genome. Unsurprisingly, soft sweeps are also much more difficult to recognize than hard sweeps because their genomic patterns are intermediate. Additionally, the identification of selective sweeps, hard or soft, is further complicated by the possibility that neutral loci linked to either soft or hard sweeps may produce a false signature similar to that of a sweep (Schrider et al. 2015; Kern and Schrider 2018).

With these challenges in mind, a considerable amount of work has been dedicated to both developing robust methods to identify selective sweeps and also understanding the evolutionary parameters that determine hard or soft sweeps. Mutation-limited scenarios are expected to exclusively produce hard sweeps because beneficial alleles rarely occur (Hermisson and Pennings 2017). Thus, the most important parameter for estimating the likelihood of hard vs soft sweeps is the population-scaled mutation rate: *Θ* = 4*N*_*e*_*μ*, where *N*_*e*_ is the effective population size and *μ* is the mutation rate. However, this single parameter can vary widely depending on the advantage of the beneficial allele, the effective population size, the size of the mutational target, and the timescale for adaptation (Messer and Petrov 2013; Hermisson and Pennings 2017). Therefore, adaptation across the genome for a given population can be simultaneously mutation-limited and non-mutation-limited (B.A. Wilson et al. 2014). While it has become clear that most populations will likely exhibit a mosaic of hard and soft sweeps (Hermisson and Pennings 2017), additional data on sweep type frequencies in various species are sorely needed to better tease apart which parameters may determine each of those frequencies.

Both species of the *Pan* genus represent important evolutionary models due to their phylogenetic proximity to humans. *Homo* and *Pan* diverged ~ 5 to 7 Ma (Sarich and Wilson 1967; Bradley 2008; Scally et al. 2012; Besenbacher et al. 2019) and the most recent estimates for the divergence of bonobos and chimpanzees range between 1 and 2 Ma (Prüfer et al. 2012; de Manuel et al. 2016). Four extant chimpanzee subspecies evolved from a chimpanzee common ancestor that split ~ 600 Ka with both subsequent lineages further splitting: one ~ 250 Ka and the other ~ 160 Ka (de Manuel et al. 2016). These two species exhibit stark differences in aspects of their morphology, physiology, behavior, and ecology (Susman 1984; Goodall 1986; Wrangham 1986; Kano 1992; White 1996; Furuichi 2011; Nishida 2011; Stumpf 2011; Behringer et al. 2014; Turley and Frost 2014; M.L. Wilson et al. 2014). Many of these distinguishing traits are inferred to have occurred shortly after divergence, while much less is known about recent evolutionary processes in these lineages.

Understanding recent positive selection in *Pan* is intriguing because of the dynamic physical and social environments in which they evolved. Climatic variation across Africa is well-documented for the Pleistocene and has been proposed to drive the evolution of early *Homo* (Potts 1998; Antón et al. 2014), and such variation probably impacted other taxa throughout the Pleistocene, including the genus *Pan*. Chimpanzee populations living in more stable environments that were closer to Pleistocene refugia were recently described to exhibit less behavioral diversity than chimpanzees living in more seasonal habitats that are more distant to forest refugia (Kalan et al. 2020). While the formation of these refugia may have resulted in periods of habitat stability for some bonobo and chimpanzee populations during glacial periods (Takemoto et al. 2017; Barratt et al. 2020), climatic fluctuations throughout the Pleistocene likely affected both the physical environment—via changes in habitat structure and type—and the social environment—via changes in the frequency of dispersal and intergroup encounters. Further, evidence of admixture within extant and between extant and extinct members of the *Pan* genus adds even more variation to the social environments in which these apes evolved (Hey 2010; Wegmann and Excoffier 2010; de Manuel et al. 2016; Kuhlwilm et al. 2019). A dynamic environment may result in selection for multiple existing alleles, resulting in a greater frequency of soft sweeps than in a more stable environment where one would expect a greater frequency of hard sweeps.

In this study, we apply a recently developed supervised machine-learning approach to population-level genomic data for bonobos (*Pan paniscus*) and chimpanzees (*Pan troglodytes*) to assess the extent of different completed sweep types in these species. While a few studies have examined recent positive selection in bonobos and chimpanzees (e.g., Cagan et al. 2016; Han et al. 2019; Schmidt et al. 2019; Kovalaskas et al. 2020; Nye et al. 2020), the role of hard and soft sweeps in shaping their adaptations is currently unknown. We sought to categorize genomic regions as subject to recent hard or soft sweeps, as linked to recent hard or soft selective sweeps, or as evolving neutrally. Data from simulations have predicted that hard sweeps would be common in humans because of our overall low mutation rate (Hermisson and Pennings 2017).

Under this “mutation limitation hypothesis” and given the similarity in mutation rate between *Homo* and *Pan*, one could predict that bonobos and chimpanzees should also exhibit a high degree of hard sweeps. However, hard sweeps have been thought and observed to be quite rare in recent human evolution (Hernandez et al. 2011; Schrider and Kern 2017), although this perspective is debated (Jensen 2014; Harris et al. 2018). This could be explained by several non-mutually exclusive alternatives including demographic effects. Larger populations can have more standing variation for selection to act on (Hermisson and Pennings 2005) which may result in more soft sweeps, whereas bottlenecks can result in drift and thus potentially more hard sweeps if intermediate frequency haplotypes are lost (B.A. Wilson et al. 2014). For example, humans have experienced recent demographic changes (e.g., Schiffels and Durbin 2014), including a bottleneck upon leaving Africa (e.g., Henn et al. 2012). Indeed, Schrider and Kern (2017) found that hard sweeps were more frequent in non-African than African populations. Chimpanzees and bonobos have also experienced recent demographic changes, including in effective population size, within the time frame (< 200 Ka) for selective sweeps, based on PSMC analyses (Prado-Martinez et al. 2013; de Manuel et al. 2016). Three of the five lineages appear to have declined, whereas the other two have increased and then decreased. Under such changes in population size, the strength of selection plays a strong role in the likelihood of soft sweeps (B.A. Wilson et al. 2014). We therefore predicted that we would observe a higher frequency of soft sweeps in *Pan*, but that lineage-specific population histories might affect the degree to which soft sweeps dominate.

## Methods

### Genomic Data

We retrieved raw short read data on bonobos and all four chimpanzee subspecies from the Great Ape Genome Project (GAGP) (Prado-Martinez et al. 2013). This dataset contained high coverage genomes (Figures S1, S2) from 13 bonobos (*P. paniscus*), 18 central chimpanzees (*P. troglodytes troglodytes*), 19 eastern chimpanzees (*P. t. schweinfurthii*), 10 Nigeria-Cameroon chimpanzees (*P. t. ellioti*), and 11 western chimpanzees (*P. t. verus*) (File S1).

### Read Mapping and Variant Calling

Initial quality assessments in fastqc (Andrews 2010) and multiqc (Ewels et al. 2016) indicated a number of quality issues, including failed runs, problematic tiles, and substantial variation in base quality. We removed adapters and trimmed all reads for quality with BBduk (https://sourceforge.net/projects/bbmap/). For trimming, we used the parameters “ktrim=r k=21 mink=11 hdist=2 qtrim=rl trimq=15 minlen=50 maq=20” for all reads and added “tpo and tpe” for paired reads.

We used XYalign (Webster et al. 2019) to create versions of the chimpanzee reference genome, panTro6 (Kronenberg et al. 2018), for male- and female-specific mapping. Specifically, the version of the reference for female mapping has the Y chromosome completely masked, as its presence can lead to mismapping (Webster et al. 2019). We then mapped reads with BWA MEM (Li, unpublished data) and used SAMtools (Li et al. 2009) to fix mate pairs, sort BAM files, merge BAM files per individual, and index BAM files. We use Picard (Broad Institute 2018) to mark duplicates with default parameters, before calculating BAM statistics with SAMtools. We next measured depth of coverage with mosdepth (Pedersen and Quinlan 2018), removing duplicates and reads with a mapping quality less than 30 for calculations.

Visualizations for coverage and demography (see Generation of Simulated Chromosomes below) were created in R, version 3.5.2 (R Core Team 2020), using ‘ggplot2’ (Wickham 2016).

We used GATK4 (Poplin et al. 2018) for joint variant calling across all samples. We used default settings for all steps—HaplotypeCaller, CombineGVCFs, and GenotypeGVCFs—with three exceptions. First, we turned off physical phasing for computational efficiency and downstream VCF compatibility with filtering tools. Second, because multiple samples in this dataset suffer from contamination from other samples both within and across taxa (Prado-Martinez et al. 2013), we employed a contamination filter to randomly remove 10% of reads during variant calling. This should have the effect of reducing confidence in contaminant alleles. Finally, we output non-variant sites to allow equivalent filtering of all sites in the genome and more accurate assessments of callability.

The above quality control, assembly, and variant calling steps are all contained in an automated Snakemake (Köster and Rahmann 2012) available on Github (https://github.com/thw17/Pan_reassembly). The repository also contains a Conda environment with all software versions and origins, most of which are available through Bioconda (Grüning et al. 2018).

### Variant Filtration and Genome Accessibility

We considered only autosomes for this analysis as the X and Y chromosome violate many of the assumptions for the following methods (Webster and Wilson Sayres 2016). We also excluded unlocalized scaffolds (N = 4), unplaced contigs (N = 4,316), and the mitochondrial genome from any downstream analyses. Additional filtration steps were completed using bcftools (Li 2011); command line inputs are provided in parentheses. Given our focus on selective sweeps, we only included single nucleotide variants (SNVs) (“-v snps”) that were biallelic (“-m2 -M2”). On a per sample basis within each site, we marked genotypes where sample read depth was less than 10 and/or genotype quality was less than 30 as uncalled (“-S . -i FMT/DP ≥ 10 && FMT/GT ≥ 30”). To ensure that missing data did not bias our results, we further excluded any sites where less than ~ 80% of individuals (N = 56) were confidently genotyped (“AN ≥ 112”). We also removed any positions that were monomorphic for either the reference or alternate allele (“AC > 0 && AC ≠ AN”). These filtrations steps yielded 41,869,892 SNVs for our downstream analyses (Table S1).

We considered sites in our sample with low to no coverage to be ‘inaccessible’ in the reference genome. Using the output of mosdepth (see Read Mapping and Variant Calling above), we identified and filtered sites exhibiting low coverage as defined above. We used the ‘maskfasta’ function in bedtools (Quinlan and Hall 2010) to mark these sites (N) in the pantro6 FASTA, featuring only the autosomes, for use in downstream analyses. This resulted in 86.3% of the assembled autosomes as accessible (File S2).

### Generation of Simulated Chromosomes

We used the software ‘discoal’ to generate simulated chromosomes on which we trained a classifier per lineage (Kern and Schrider 2016). We generated a matching number of simulated haploid chromosomes for the sample size of each *Pan* lineage (i.e., 26 chromosomes for 13 *P. paniscus*, 20 chromosomes for 10 *P. t. ellioti*, etc.). Simulated chromosomes were set to 1.1 Mb in length and divided into 0.1 Mb subwindows for a total of 11 subwindows. These simulations included a population-scaled mutation rate (4*NμL*), where *N* is the effective population size, *μ* is the per base pair per generation mutation rate, and *L* is the length of the simulated chromosome. We used the median of the previously reported effective population size range per lineage (Prado-Martinez et al. 2013). As estimates of genome-wide mutation rates vary considerably and are complicated in that mutation rates vary across individual genomes, we based our parameter on a mutation rate of 1.6 × 10^−8^, which falls between estimates from genome-wide data and phylogenetic estimates (Narasimhan et al. 2017). We introduced some variation in this rate by setting a lower and upper-bound to 1.5 and 1.7 × 10^−8^ and sampled a new mutation rate per simulation drawing from this uniform prior. All simulations also included a population-scaled recombination rate (4*NrL*), where *r* is the recombination rate per base pair per generation, again calculated from the median effective population size for each lineage from Prado-Martinez et al. (2013) and a recombination rate drawn from a uniform prior of 1.1 − 1.3 × 10^−8^, based on the mean genome-wide rate (1.2 × 10^−8^) reported for bonobos, chimpanzees, and gorillas (Stevison et al. 2015). Recent results from a different selective sweep classifier, Trendsetter, suggest that including a range of recombination rates is important to reducing misclassification (Mughal and DeGiorgio 2019). We note that while some of the estimated recombination rates in bonobos and chimpanzees are beyond the uniform distribution used in our simulations, many of these values are the high rates present in the telomeres, regions that generally exhibit lower or no coverage and thus will be largely if not entirely masked from this analysis (see Variant Filtration and Genome Accessibility above). We also included a demographic string reflecting approximate changes in population size for each lineage between ~ 0.05 and 2 Ma. Changes in population size were set in units of 4N_0_ generations, N_0_ was set to the approximate median effective population size from (Prado-Martinez et al. 2013) and we used a generation time of 25 years (Langergraber et al. 2012). Population size changes for this time period were drawn from a previous PSMC analysis (de Manuel et al. 2016) (Figure S3). While this is only one study from which to draw demographic information and reconstructions of *Pan* demography vary widely across studies, the downstream program used to classify genomic windows, diploS/HIC, is robust to demographic misspecification (Kern and Schrider 2018). We generated 2 × 10^3^ simulations using these parameters as a set of simulations under neutral evolution per lineage.

Hard and soft selective sweeps were simulated with all of the aforementioned parameters and using a uniform prior of population-scaled selection coefficients (α = 2*Ns*) derived from each lineage’s median effective population size (Prado-Martinez et al. 2013) and moderately weak to moderately strong selection coefficients between 0.02 and 0.05. Sweeps also included a parameter (τ) for the time to fixation of the beneficial allele over a uniform range in units of 4N generations. This value ranged from 0 to 0.001 for all lineages. Linked-hard and linked-soft sweeps were generated by placing the selected site at the center of each of the 10 subwindows flanking the center (6^th^) subwindow. Additionally, we included a uniform prior on the frequency at which a mutation is segregating at the time it becomes beneficial for soft and linked-soft sweeps, setting this range from 0 to 0.2. We generated 1 × 10^3^ simulations per subwindow for linked-hard and linked-soft sweeps (N = 10) and 2 × 10^3^ simulations for hard and soft sweeps. This resulted in a total of 2 × 10^3^ hard, 1 × 10^4^ hard-linked, 2 × 10^3^ soft, and 1 × 10^4^ soft-linked simulated sweeps. Parameters for these simulations are presented in File S3.

### Calculation of Simulation Feature Vectors and Classifier Training

We calculated feature vectors from these simulated chromosomes using the ‘fvecSim’ function in the program diploS/HIC (Kern and Schrider 2018). Briefly, diploS/HIC calculates 12 summary statistics for all 11 subwindows: *π*, Watterson’s *θ*, Tajima’s *D*, the variance, skew, and kurtosis of genotype distance (*g*_*kl*_), the number of multilocus genotypes, *J*_1_, *J*_12_, *J*_2_/*J*_1_, unphased *Z*_*ns*_, and the maximum value of unphased ω. Collectively, these summary statistics capture information about the site frequency spectrum (SFS), haplotype structure, and linkage disequilibrium (LD). diploS/HIC uses a convolutional neural network (CNN) to capture essential aspects of a feature (the feature vector) by sliding a receptive field over the image to compute dot product between the original filter and the convolutional filter. In diploS/HIC, the CNN uses three branches of a CNN, of which each has two dimensional convolutional layers with ReLu activations followed by max pooling. This is followed by a dropout layer to control for model overfitting. Outputs from all three units are fed into two fully connected dense layers, which also use dropout layers, before arriving at a softmax activation that outputs the probability for each categorical class (hard, hard-linked, neutral, soft-linked, or soft). Complete details for this procedure can be found in Kern and Schrider (2018).

When calculating feature vectors for the simulated chromosomes, we used the optional arguments for the ‘fvecSim’ function to mask each simulation with 110,000 bp segment randomly drawn from our masked FASTA where > 0.25 of SNVs in a subwindow were accessible (i.e., not marked by Ns). This enabled us to train our classifiers on simulated data featuring the same patterns of inaccessible genomic regions that the classifier would encounter in the empirical data.

We created a balanced set with equal representation (2 × 10^3^) of all five classes via sampling without replacement in which to train the classifier using diploS/HIC’s ‘makeTrainingSets’ function. These were divided into 8,000 training examples, 1,000 validation examples, and 1,000 testing examples to test the accuracy of the classifier via the ‘train’ function in diploS/HIC. We built ten classifiers per lineage and selected the one with the highest accuracy to apply to the empirical data (File S4).

A second, independent set of simulated chromosomes was generated per lineage using the same parameters. We then calculated feature vectors and created another balanced training set with 2 × 10^3^ chromosomes per class (hard, linked-hard, neutral, linked-soft, and soft). We used diploS/HIC’s ‘predict’ function by applying each trained classifier to all five classes separately per lineage. In other words, we ran each classifier on 2000 simulated hard sweeps, 2000 simulated linked-hard sweeps, 2000 simulated neutral regions, 2000 simulated linked-soft sweeps, and 2000 simulated soft sweeps and for each lineage. We used a binary classification scheme, where the identification of a sweep (hard or soft) was considered to be positive and linked or neutral regions were negative, to assess the true positive rate, false positive rate, and obtain a second estimate of accuracy for each trained classifier (Tables S2 − S5). We also calculated class-specific accuracy, by summing the number of instances per lineage where the predicted class matched the simulated class divided by the total (1 × 10^4^) (Tables S2 - S5).

### Empirical Data Feature Vectors and Prediction

Upon achieving > 0.8 accuracy, each trained classifier was applied to its respective *Pan* lineage. Each autosome was analyzed separately and feature vectors calculated using diploS/HIC’s ‘fvecVcf’ function. We supplied this function with the masked FASTA for that chromosome and discarded windows where any subwindow had < 0.25 unmasked sites following Schrider and Kern (2017) (File S5). This step reduces the potential effect of the number of SNVs in a given window on sweep classification. Finally, the trained classifier was applied to the feature vector files using the ‘predict’ function.

### Sweep Identification, Potential Target Genes, and Gene Ontology

As diploS/HIC outputs the probability for each sweep class, we first report the class inferred to be the most likely. However, as the difference between the most likely class and the next most likely may be small, we further report windows where the sweep class probability is > 0.5, > 0.75, and > 0.9 (File S6). We also examined our data for spatial patterns. Windows classified as immediately abutting other windows with the same sweep type for hard and soft sweeps were considered to be a single sweep. Unique sweep windows and those shared between two or more lineages were visualized using UpSet plots (Lex et al. 2014) in R (R Core Team 2020).

We examined what genes lie in the windows identified as being subject to a recent selective sweep by extracting the genomic coordinates of all autosomal coding regions for the longest transcript per gene (N = 20,119 genes) in the panTro6 genome via the panTro6 gff (retrieved from: https://www.ncbi.nlm.nih.gov/genome/202?genome_assembly_id=380228). We used the bedtools ‘intersect’ function (Quinlan and Hall 2010) to identify overlap between coding regions and candidate sweep windows after converting both CDS and sweep window coordinates to 0-start, half-open format. As some coding sequences may have been masked (see Variant Filtration and Genome Accessibility above), we extracted FASTAs for each coding sequence using bedtools ‘getfasta’ function (Quinlan and Hall 2010) and used a custom R script to calculate the percent of each gene that was masked. Overall, 66.2% of all coding sequence was unmasked. We excluded listing genes for candidate sweep regions if > 50% of the total coding sequence per gene was masked. Thus, we considered 13,228 genes as potential targets for selective sweeps (File S7).

We investigated the enrichment of particular pathways by performing a gene ontology analysis using the Functional Annotation Tool in DAVID (Huang et al. 2008; Huang et al. 2009). We used the custom background described above (genes whose total coding sequence was > 50% unmasked) rather than all pantro6 genes to ensure our analysis was not underpowered.

DAVID does not allow for official gene symbols to be used in a background list, so we converted gene symbols to Entrez gene IDs. As not all gene symbols have a corresponding Entrez gene ID, we removed genes for which there was no Entrez gene ID (N = 98 in background list). We collated genes for both hard and soft sweeps into a single input per lineage. We evaluated statistical significance for biological process gene ontology terms via p-values adjusted using the Benjamini-Hochberg method (Benjamini and Hochberg 1995).

Scripts for all data analyses are available on Github (https://github.com/brandcm/Pan_Selective_Sweeps).

## Results

We generated four classifiers that reached an acceptable level of accuracy for bonobos (*P. paniscus*), central chimpanzees (*P. t. troglodytes*), eastern chimpanzees (*P. t. schweinfurthii*), and Nigeria-Cameroon (*P. t. ellioti*) chimpanzees. These classifiers ranged in accuracy from 85.6% (Nigeria-Cameroonian chimpanzees) to 93.9% (central chimpanzees) (File S4). We could not produce a sufficiently accurate classifier using realistic parameters for western chimpanzees (*P. t. verus*); therefore, they were excluded from downstream analyses. Our trained classifiers had considerable statistical power (1 - false positives) ranging from 96.6 to 99.2% and a low false positive rate (false positives / false positives + true negatives) that ranged from 1.4 to 4.3% across all four classifiers (Tables S2 - S5). When considered separately—i.e., true positives only included one sweep type (hard or soft) rather than both—we had greater power to detect hard sweeps than soft sweeps, averaging 99% and 96.9% across lineages, respectively (Tables S2 - S5). Accuracy (true positives + true negatives / total) for identifying sweep regions vs non-sweep regions ranged from 94.1 to 98.3% while a second estimate (in addition to the first accuracy estimate that resulted from the construction of the classifiers) of class-specific accuracy ranged from 81.6 to 92.1% (Tables S2 - S5).

We classified ~ 91.6% of the assembled autosomes in each lineage (Table 1, File S8), even after masking for inaccessible regions and excluding windows with few SNVs. We found that soft sweeps were abundant in all four lineages, accounting for > 73% of all individual sweeps, whereas hard sweeps were relatively rare (Table 1, File S8). This pattern held true even when more stringent posterior probabilities were applied to consider a region a sweep and at least 30% of hard sweep windows and 76% of soft sweep windows were called with 50% or greater posterior probability (File S6). Genomic regions linked to sweeps were also quite pervasive in all four lineages (Table 1); particularly among eastern chimpanzees, where roughly 86% of the genome was classified as linked to selective sweeps.

**Table 1.** Selective sweep summary per population.

| Lineage | Number / Percent of Windows per Class Type |  |  |  |  |  | Number and Percent of Sweep Type |  |  |
| --- | --- | --- | --- | --- | --- | --- | --- | --- | --- |
|  | Hard | Linked-hard | Neutral | Linked-soft | Soft | Total | Hard | Soft | Total |
| *P. paniscus* | 85<br>(0.4%) | 1,576<br>(6.5%) | 7,488<br>(30.8%) | 13,168<br>(54.1%) | 2,002<br>(8.2%) | 24,319 | 81<br>(4.9%) | 1,585<br>(95.1%) | 1,666 |
| *P. t. ellioti* | 573<br>(2.4%) | 6,358<br>(26.1%) | 1,389<br>(5.7%) | 14,498<br>(59.6%) | 1,505<br>(6.2%) | 24,323 | 488<br>(26.9%) | 1,323<br>(73.1%) | 1,811 |
| *P. t. schweinfurthii* | 32<br>(0.1%) | 696<br>(2.9%) | 1,835<br>(7.5%) | 20,179<br>(83.0%) | 1,581<br>(6.5%) | 24,323 | 32<br>(2.3%) | 1,376<br>(97.7%) | 1,408 |
| *P. t. troglodytes* | 224<br>(0.9%) | 1,746<br>(7.2%) | 5,483<br>(22.5%) | 15,121<br>(62.2%) | 1,749<br>(7.2%) | 24,323 | 184<br>(10.6%) | 1,557<br>(89.4%) | 1,741 |

We examined overlap in windows classified as either a hard or soft sweep across lineages, which may reflect either ancestral or parallel adaptation. Most hard sweep windows were unique to each lineage; however, we did find some shared windows across lineages (Figure 1). Central and Nigeria chimpanzees shared the highest number of sweep windows (N = 33) but when weighted by the total possible number of windows, the highest overlap for hard sweeps was between eastern and Nigeria chimpanzees (7/32 or ~ 0.21). No hard sweeps windows were shared across all lineages. Like hard sweeps, most soft sweep windows were also unique to each lineage (Figure 2). Among pairs of lineages there was remarkable consistency in the number of shared windows (N = 111-147), even when the total possible number of shared windows is considered. One exception is eastern and central chimpanzees who shared nearly twice the number of soft sweep windows (N = 267). The highest number of shared soft sweep windows between three lineages occurred in the three chimpanzee subspecies (N = 80). Only 19 windows were shared across all four lineages.

**Figure 1.**
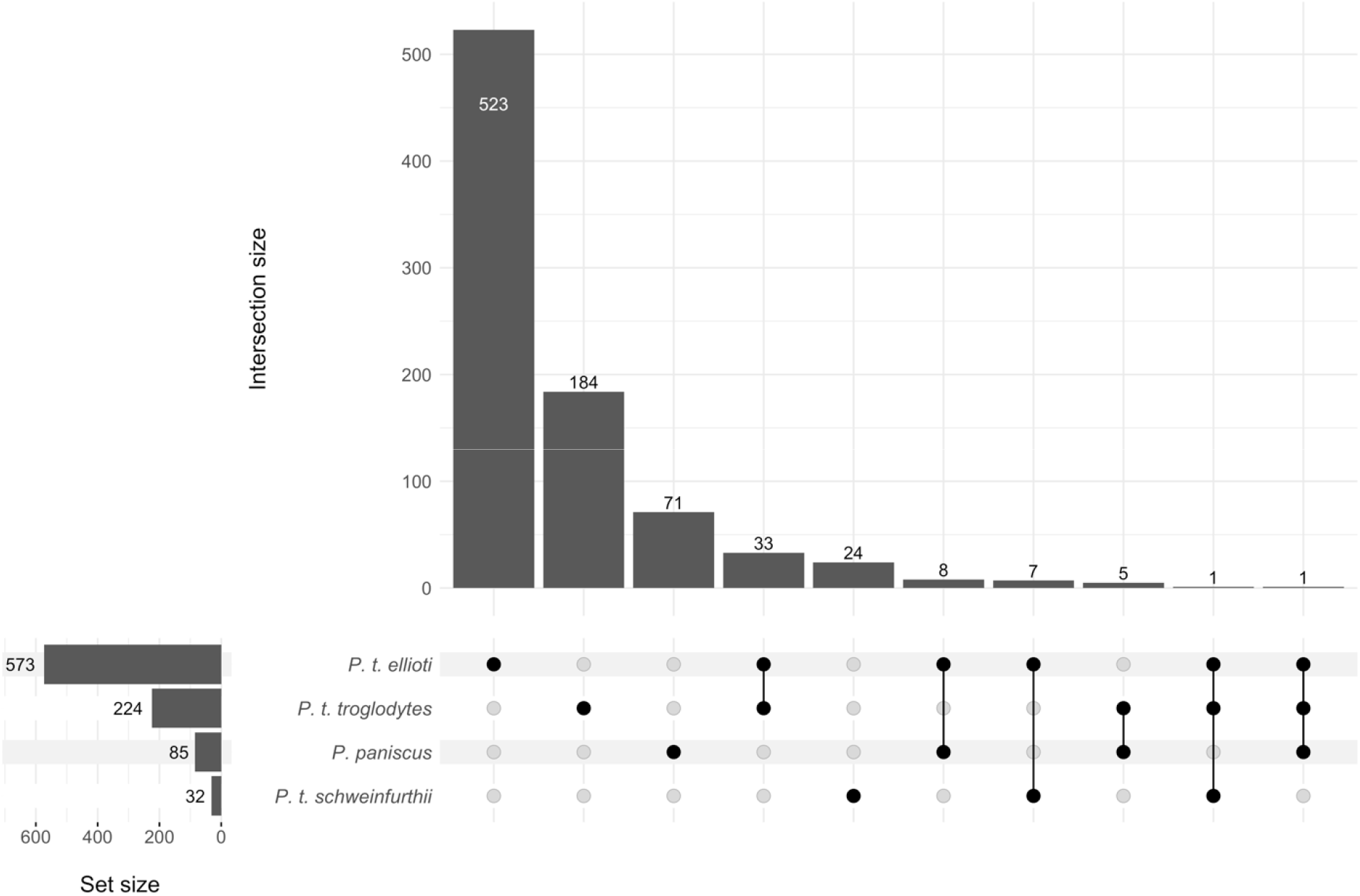
Unique and shared hard sweep windows. The frequency of windows shared by two or more lineages should be considered relative to the total possible number of shared windows (i.e., the set size of the lineage with the smallest set size).

**Figure 2.**
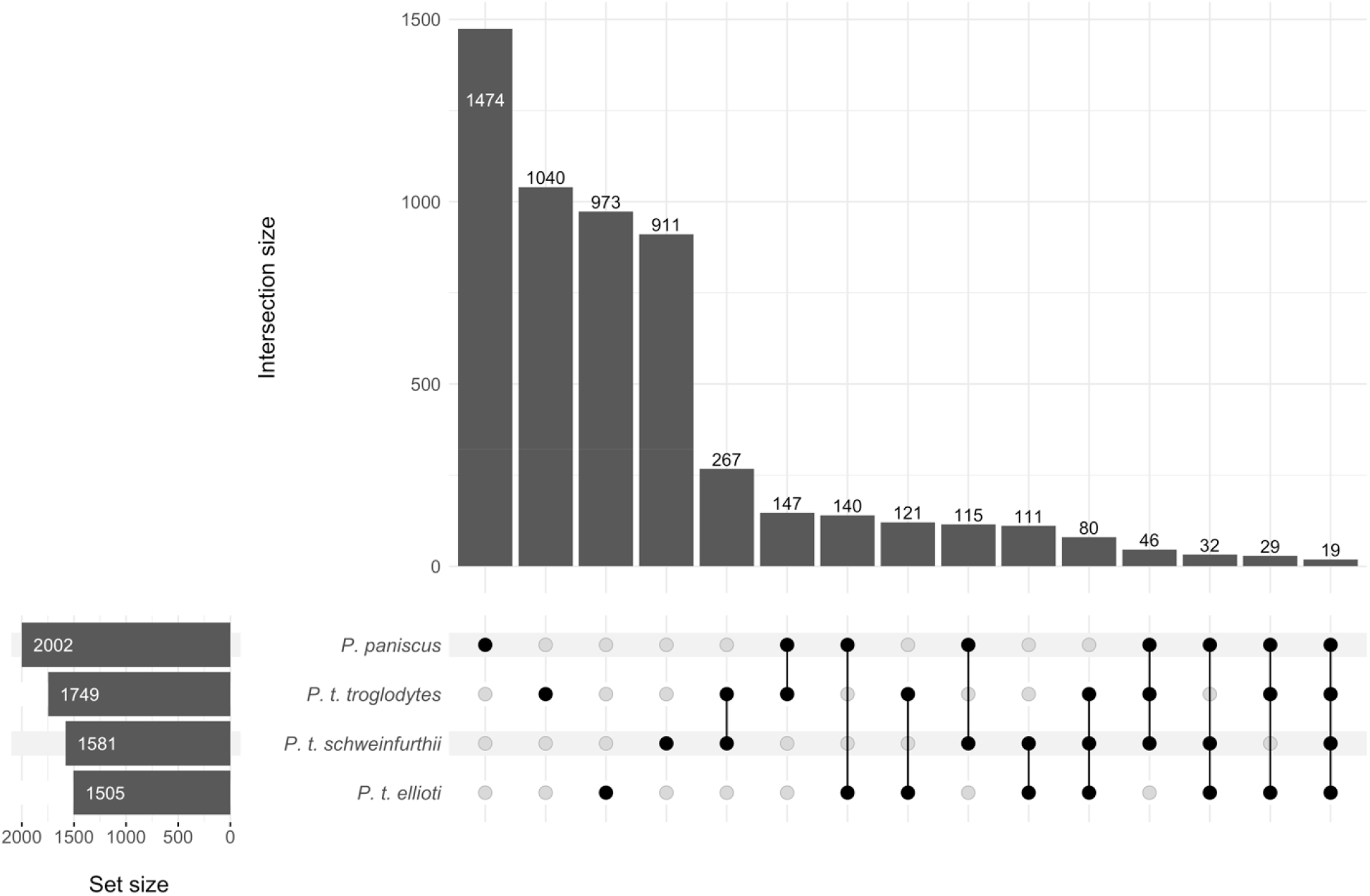
Unique and shared soft sweep windows. The frequency of windows shared by two or more lineages should be considered relative to the total possible number of shared windows (i.e., the set size of the lineage with the smallest set size).

After excluding genes that were > 50% masked, we identified 1,671 candidate genes in bonobo hard and soft sweeps, 1,761 genes in central chimpanzee sweeps, 1,372 genes in eastern chimpanzee sweeps, and 1,844 genes in Nigeria-Cameroonian chimpanzee sweeps (File S9). After correcting for multiple testing, across all lineages, we identified only two significantly enriched pathways in central chimpanzees: nervous system development and central nervous system development (File S10).

## Discussion

Our study contributes to the emerging picture of recent evolution in *Pan* and adaptation more broadly. Contrary to the predictions of a mutation-limitation hypothesis, yet concordant with recent results for humans (e.g., Hernandez et al. 2011; Schrider and Kern 2017) and flies (Garud et al. 2015), we find soft sweeps to overwhelmingly predominate regions of the genome experiencing selective sweeps in both bonobos and the three chimpanzee subspecies we could analyze. These results confirm the prediction from Schmidt et al. (2019) who speculated that soft sweeps played a major role in the evolution of eastern and central chimpanzees. Those authors also posit that hard sweeps should be more frequent in western chimpanzees relative to other subspecies because of their low effective population size. While western chimpanzees are estimated to have the lowest effective population size, it is estimated to be only slightly lower than that of bonobos for which we found a high number (95.1%) of soft sweeps (e.g., Prado-Martinez et al. 2013; de Manuel et al. 2016). It is curious that Nigeria-Cameroon chimpanzees exhibit the most hard sweeps in this analysis. While this could be the result of a multitude of factors, it is particularly curious because this lineage has experienced a rather stable effective population size in recent evolutionary time as estimated by PSMC (Prado-Martinez et al. 2013; de Manuel et al. 2016), whereas a scenario with dramatic population decline would be expected to “harden” soft sweeps as haplotypes are stochastically lost, resulting in more hard sweeps (B.A. Wilson et al. 2014).

Our analysis of shared hard and soft sweeps found that most sweeps of both types were unique to each lineage. However, there was a high number of hard sweep windows shared between central and Nigeria-Cameroon chimpanzees as well as between eastern and Nigeria-Cameroon chimpanzees when the total possible number of shared sweeps was considered. Further, there were nearly twice the number of shared soft sweep windows shared between eastern and central chimpanzees. These results are similar to other recent findings (Nye et al. 2020). It is impossible to discern whether or not the overlap in hard sweeps between central and Nigeria-Cameroon chimpanzees and the overlap in soft sweeps for eastern and central chimpanzees is the result of shared ancestry and/or similar environmental conditions because both pairs of lineages share a geographic boundary: the Ubangi river for eastern and central chimpanzees and Sanaga river for central and Nigeria-Cameroon chimpanzees. The overlap in hard sweeps between eastern and Nigeria-Cameroon chimpanzees is more puzzling because they are not sister taxa and share a common ancestor ~ 600 Ka. Therefore, parallel adaptation via similar physical and/or social environments may serve as a more likely hypothesis. While the lowest in overall frequency, we also identified a number of soft sweep windows that were shared across three lineages as well as 19 windows that occurred in all four. Future work should further investigate these shared sweep windows.

As mentioned above, soft sweeps are not exclusively the result of selection on standing genetic variation (Pennings and Hermisson 2006a; Pennings and Hermisson 2006b). However, given the mutation rates estimated for bonobos and chimpanzees, it appears unlikely that recurrent *de novo* mutations explain the majority of these soft sweeps. We did not explicitly model for different types of soft sweeps in our analysis. However, while soft sweeps from standing genetic variation and *de novo* mutations may exhibit similar genomic signatures, the hypothesis that these processes result in similar genomic signatures must be tested before any additional conclusions are drawn. Hartfield and Bataillon (2020) recently suggested differences in diversity (as measured by *π*) at the selected locus may be used to differentiate soft sweep types, although this may be more difficult to accomplish in outcrossing species. Nonetheless, our results reveal a major role of standing genetic variation, and thus changes in the physical and social environment, in driving recent adaptations in *Pan*.

A few recent studies have considered the impact of effective population size on adaptive evolution in the great apes (Cagan et al. 2016; Nam et al. 2017). Theory predicts that the rate of adaptive evolution should be positively correlated with effective population size when *N*_*e*_*s* is >> 1 (Gossmann et al. 2012). Both Cagan et al. (2016) and Nam et al. (2017) found a positive association between effective population size and the rate of adaptive evolution, measured by proportion of adaptive substitutions and the number of selective sweeps, respectively. However, we observed no clear linear relationship between the number of sweeps (hard, soft, or both) estimated from this analysis and the estimated effective population sizes for these four lineages (see File S3 for population sizes). This descriptive result should be considered cautiously because of the limited number of lineages analyzed here and the potential confounding effect of phylogeny. It is possible that this relationship may not be driven by the number of sweeps, but rather the strength of sweeps a population experiences (Nam et al. 2017). Estimates of selection strength are generally lacking for the great apes so this relationship remains a question for further study.

In addition to characterizing broad patterns in the genomic landscape for bonobos and chimpanzees, the results of this study also highlight thousands of candidate regions and genes for further analysis. We also find additional support for previous selection candidates. For example, disease has been long thought to shape evolution in primates (Nakajima et al. 2008; van der Lee et al. 2017). The potential for disease transmission between non-human primates and humans has also prompted much research, particularly focusing on the genomic underpinnings of host responses to lentiviruses, which include HIV and SIV (Gao et al. 1999; Van Heuverswyn et al. 2006; Compton et al. 2013; Nakano et al. 2020). Cagan and colleagues (2016) found evidence of recent positive selection within *IDO2*, a T-cell regulatory gene, among all four-chimpanzee subspecies and bonobos. We identified a candidate soft sweep region for eastern chimpanzees that overlaps this gene. However, this window had one of the lowest posterior probabilities in this lineage (49.7%) and there was a nearly equally high probability that this window was linked to a soft sweep (43.8%). Clearly, additional work is needed to understand the potential role of *IDO2* in *Pan* evolution. Schmidt et al. (2019) recently described three chemokine receptor genes—*CCR3, CCR9*, and *CXCR6—*had a significant number of highly differentiated SNVs in central chimpanzees. We could evaluate all three of these genes in our analysis but only one fell within a candidate sweep window: *CXCR6*. The window containing this gene was confidently called as a soft sweep with a posterior probability of 85.5%. It is not known as to whether or not SIV_cpz_ uses *CXCR6* to enter chimpanzee host cells (Wetzel et al. 2018). However, multiple lines of evidence for selection either at this locus or within the window overlapping this gene prompt a closer examination of this genomic region. Finally, *TRIM5* fell within a hard sweep window in central chimpanzees. *TRIM5* is a well-known retrovirus restriction factor that appears subject to ancient, multi-episodic positive selection in primates (Sawyer et al. 2005).

Recent attention has focused on admixture between lineages in the genus *Pan* and the potential adaptiveness of introgressed genomic elements. de Manuel and colleagues (2016) identified 221 genes that fell within putatively introgressed elements in central chimpanzees from admixture with bonobos. Some of this admixture is estimated to occur < 200 Ka, thus within the timeframe that the present analysis can detect selective sweeps. While we could not evaluate six of these 221 genes, five fell within candidate sweep regions in central chimpanzees from our study: *CDK8, EIF4E3, GRID2, PTPRM*, and *TRIM5*. As described above, *TRIM5* was unique to central chimpanzees. We found *CDK8* in sweep windows for bonobos, eastern chimpanzees, and Nigeria-Cameroon chimpanzees. In humans, *CDK8* mutations have been associated with multiple phenotypic effects including hypotonia, behavioral disorders, and facial dysmorphism (Calpena et al. 2019). We also identified *EIF4E3* in candidate sweeps for bonobos whereas *GRID2* and *PTPRM* were found in eastern chimpanzees. *EIF4E3* is a translation initiation factor (Osborne et al. 2013) while *PTPRM* is a member of the protein phosphatase family (PTP) and has multiple functions including cell proliferation and differentiation (Sun et al. 2012). *GRID2* generates ionotropic glutamate receptors and mutations have been associated with abnormalities of the cerebellum (Lalouette et al. 1998).

The gene ontology analysis produced only two statistically significant terms, nervous system development and central nervous system development, for a single *Pan* lineage: central chimpanzees. While cognitive and neurological differences are widely considered to differentiate bonobos and chimpanzees (e.g., Rilling et al. 2012; Stimpson et al. 2016; Staes et al. 2019), we are unaware of any studies that investigate variation among chimpanzee subspecies that may explain enrichment for nervous system and central nervous system development related genes specifically in central chimpanzees. We note that compared to other gene ontology analyses, our level of enrichment is quite low. While we excluded a large number of genes from our analysis due to poor coverage, our use of a custom background should increase, rather than decrease, statistical power.

The results from our analysis should be interpreted with some caution. First, while our classifiers achieved a high degree of accuracy, it is possible that some selective sweeps in each lineage were not detected or regions were incorrectly identified as such (Tables S2 - S5). We also note that we did not model small selection coefficients (*s* < 0.02) as we could not accurately classify sweeps under weak selection, which may be the result of the large window size (1.1 Mb) used here. One consequence may be that if weakly beneficial hard sweeps are present in the empirical data, they may have been sometimes classified as soft (Harris et al. 2018).

Nonetheless, our classifiers were overall quite good at identifying moderately selected hard and linked-hard sweeps with both at approximately 95% accuracy across all lineages. Neutral and linked-soft regions were the most difficult to recognize with neutral regions typically being classed as soft-linked when they did not appear neutral. This suggests that the neutral portion of the genome for each lineage is slightly underestimated here. Finally, some moderately selected soft sweeps were identified as hard sweeps in each of our classifiers, suggesting that some portion of identified hard sweeps in each lineage are, in fact, soft sweeps. The low false positive rates demonstrate the overall accuracy of the observed genomic patterns (i.e., the proportion of hard and soft sweeps) for these taxa. However, this point underscores the need to conduct subsequent analyses of the candidate regions and genes to confirm such the proposed mode of adaptation and investigate any functional consequences of that adaptation. In the ‘era of -omics’, the generation of candidate regions for any type of selection across populations and species appears to overwhelmingly outpace the confirmation of such patterns. Avenues of research that investigate these candidate genes in more detail are thus well poised to provide a deeper and more accurate understanding of lineage-specific adaptations.

Second, background selection, the loss of a linked neutral site from purifying selection on a deleterious allele, can potentially mimic patterns of selective sweeps and thus may impact the results of this study (Charlesworth et al. 1993). We did not explicitly model background selection in our analysis, however, evidence from simulations in various taxa demonstrate that this pattern of selection does not substantially increase the rate of false positives in selective sweep analyses (Schrider and Kern 2017; Schrider 2020). Further, Nam et al. (2017) considered the effect of background selection on genomic diversity in extant apes, including all five *Pan* lineages, and note that background selection alone does not produce the observed diversity reduction near genic regions in these lineages. While background selection may not largely affect certain selective sweep analyses, it may impact estimations of demography that are inferred using PSMC/MSMC approaches (Johri, Riall, et al. 2020; Johri, Charlesworth, et al. 2020). The demographic strings calculated from PSMC used in this analysis also broadly agree in population size shape with other demographic estimates generated using other methods (e.g., Becquet and Przeworski (2007); Hey (2010)), therefore, background selection unlikely affects the demographic models used in analysis. Yet, this issue should be strongly considered in future studies where demography is only inferred from PSMC/MSMC.

Further, sampling bias can reduce the accuracy of identifying selective sweeps. If multiple haplotypes are present in a population but only individuals sharing one haplotype are sampled, then the sweep would be classified as a hard sweep when it is a soft sweep. However, this scenario would only underestimate the degree of recent adaptation from soft sweeps. Therefore, if this sampling bias is present in this analysis, then soft sweeps may predominate recent *Pan* evolution to an even larger degree than described here. Population structure adds further complications to the classification of hard sweeps. Parallel adaptation produces multi-origin soft sweeps at the global population level that would appear to be hard in local populations, although even local samples may sometimes appear to be soft sweeps (Ralph and Coop 2010). Thus, if samples stemmed from one or few local populations then global soft sweeps may be misclassified as hard. A previous analysis estimated the geographic origin of individuals used in this analysis (de Manuel et al. 2016). These authors found that individuals from both eastern and central chimpanzee populations were sampled from multiple countries across the geographic range for both subspecies. Therefore, any hard sweeps detected in these populations are likely accurate at the subspecies level. Geographic origin could not be assessed for any of the bonobos or all of the Nigeria-Cameroon chimpanzees used in this analysis (de Manuel et al. 2016). As such, sampling or geographic bias may partially explain the high degree of hard sweeps observed in Nigeria-Cameroon chimpanzees, if they were sampled from a smaller geographic area than the other subspecies. We encourage future studies to consider this potential bias when hard sweeps are encountered in existing data and during study design.

This analysis focuses on signatures of positive selection at single loci. However, there is theoretical and empirical evidence that a number of adaptive traits have a complex, multilocus architecture (Pritchard et al. 2010; Yang et al. 2017; Bergey et al. 2018). For these polygenic traits, shifts in the physical or social environment might result in allele frequency changes at many loci, of which, according to models, few to none of which would reach fixation (Pritchard et al. 2010). This may, in part, explain why hard sweeps appear to be rare in humans and other species if it represents a dominant mode of adaptation in these taxa. Unfortunately, at this point, we lack the data and methods to investigate the extent of polygenic selection across the genome in many non-model taxa such as *Pan*. Another factor to consider is dominance. Here, we assumed advantageous alleles were codominant, however, there is evidence that dominance may influence patterns of selective sweeps when variants occur via *de novo* mutation or recurrent mutation (Hartfield and Bataillon 2020). It is also worthwhile to address that this analysis explicitly focused on modelling very recent completed selective sweeps. Another future avenue of study in these lineages is the identification of incomplete or partial sweeps using existing approaches (Ferrer-Admetlla et al. 2014; Vy and Kim 2015) as well as explicitly modelling both incomplete and complete sweeps to address potential “temporal misclassification” (Zheng and Wiehe 2019).

Finally, while our approach to identifying hard and soft sweeps is a logical first step, future work should consider sweeps within subspecies to assess population-level (i.e., local), rather than lineage-specific (i.e., global) adaptations. This is underscored by the extensive phenotypic variation among chimpanzees, particularly that of behavioral variation, which includes key characteristics that are often used to dichotomize bonobos and chimpanzees (Wilson et al. 2014). Further investigation is also clearly warranted in bonobos, whose overall phenotypic variation is likely underappreciated compared to chimpanzees (Hohmann and Fruth 2003; Sakamaki et al. 2016; Beaune et al. 2017; Wakefield et al. 2019).

## Conclusion

This study highlights the importance of changes in physical and/or social environment via soft selective sweeps in the recent evolution of our closest living relatives, chimpanzees and bonobos. Our results also yield further support for the ubiquity of soft, rather than hard, sweeps in adaptation. We contribute candidate regions and genes that may help identify unique phenotypes in each *Pan* lineage. Our findings also prompt many new questions including the estimation of selection strength coefficients and the degree of haplotypic diversity in candidate sweep regions. While our study focuses on these lineages broadly, this point also underscores the need for high-coverage genomic data collected using non-invasive methods at more local geographies.

## Acknowledgements

We thank Andy Kern for help with implementing this analysis. Hazel Byrne, Tina Lasisi, Alan Rogers, Liz Tapanes, and Andrew Zamora provided valuable comments on this manuscript. We also thank Elisabeth Goldman and Noah Simons for assistance with bioinformatics. We gratefully acknowledge Brad Sherman (NIH) who provided assistance with our gene ontology analysis. We thank Mark Allen, Mike Coleman, and Rob Yelle (University of Oregon Research and Advanced Computing Services) for their help with use of UO’s computing cluster—Talapas. Finally, we thank the Center for High Performance Computing at the University of Utah for resources and support.

## Supplements

- Main Supplemental File: Figures S1 - S3, Tables S1-S4.
- File S1. Sample information. (File name: File_S1_sample_information.xlsx)
- File S2. Genome accessibility information. (File name: File_S2_genome_accessibility.xlsx)
- File S3. Discoal parameter information. (File name: File_S3_discoal_input_summary.xlsx)
- File S4. Classifier trial information. (File name: File_S4_diploshic_classifier_summary.xlsx)
- File S5. Unmasked SNV count/fraction per window for VCF feature vectors. (File name: File_S5_fvec_vcf_unmaskedsnpcount_unmaskedfrac_summary)
- File S6. Number of hard and soft sweep windows using higher probability thresholds. (File name: File_S6_sweeptype_probability_cutoff_summary.xlsx)
- File S7. Genes included in sweep analysis (File name: File_S7_genes_to_include.xlsx)
- File S8. Sweep information. (File name: File_S8_selective_sweep_summary.xlsx)
- File S9. List of genes in hard and soft sweeps. (File name: File_S9_gene_lists.xlsx)
- File S10. Gene ontology analysis. (File name: File_S10_gene_ontology.xlsx)

