## Supplementary material for "Soft sweeps predominate recent positive selection in bonobos (*Pan paniscus*) and chimpanzees (*Pan troglodytes*)": Figures S1-S3, Tables S1-S4

Main Supplemental File.

Figure S1. Inter-individual variation in mean total genomic coverage and grouped by population. Total includes all autosomes, sex chromosomes, and the mitochondrial genome. Dashed lines represent means per population.

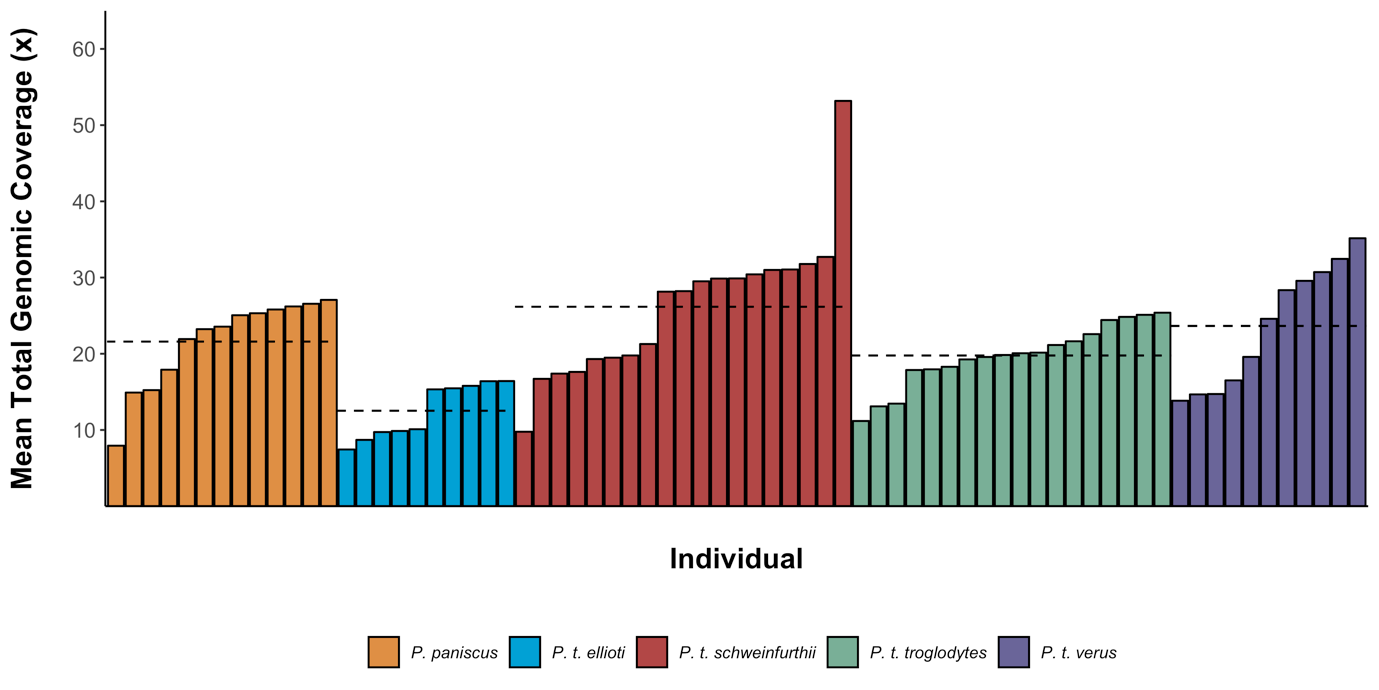

Figure S2. Inter-chromosomal variation in genomic coverage for all autosomes. Violin plots represent density at a particular coverage with the individual values plotted and jittered for each chromosome.

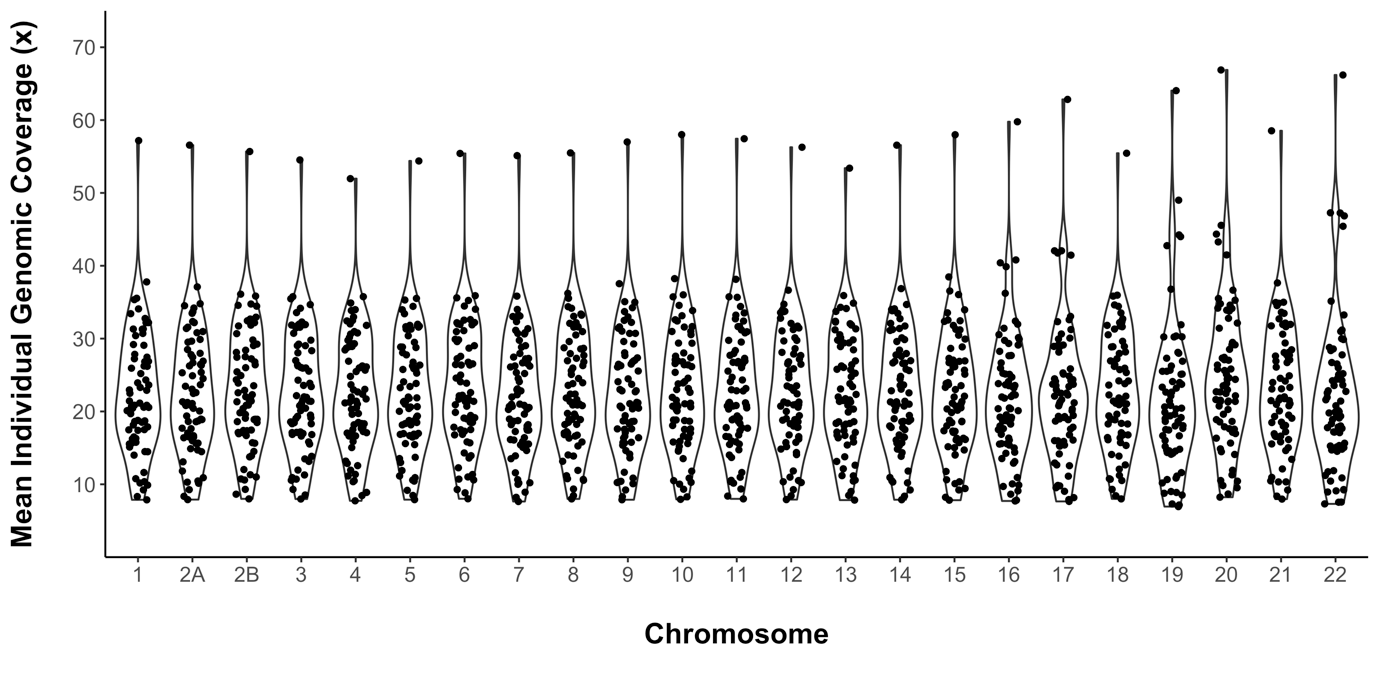

Figure S3. Plot of demographic strings used in simulated data. Demography data adapted from de Manuel et al. 2016. Data points are smoothed here using a spline function.

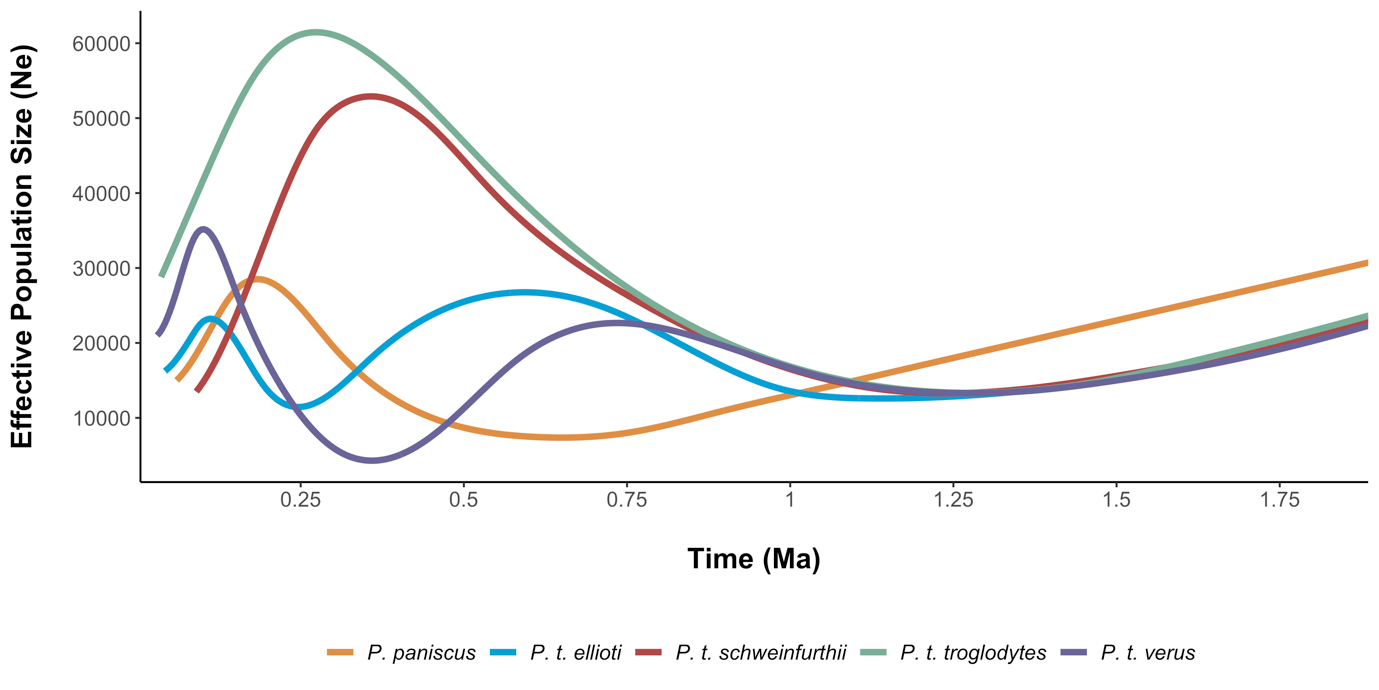

Table S1.

| chromosome | N SNVs post filtering |
| --- | --- |
| 1 | 3332313 |
| 2A | 1774559 |
| 2B | 2024437 |
| 3 | 3156661 |
| 4 | 3086706 |
| 5 | 2585495 |
| 6 | 2741379 |
| 7 | 2451758 |
| 8 | 2415460 |
| 9 | 1815711 |
| 10 | 2086543 |
| 11 | 2056912 |
| 12 | 2024301 |
| 13 | 1570832 |
| 14 | 1373336 |
| 15 | 1246426 |
| 16 | 1202275 |
| 17 | 1016097 |
| 18 | 1254126 |
| 19 | 687464 |
| 20 | 971965 |
| 21 | 560810 |
| 22 | 434326 |
| Total | 41869892 |

Table S2. Simulated vs predicted sweep type classes for *P. paniscus* classifier on a second, independent set of simulations with identical parameters. We generated 2 x 10^3^ simulations for each class. Shaded cells reflect accuracy as a binary classifier: positive if hard or soft sweep, negative if linked sweep or neutral. Green = true positive, blue = false negative, purple = false positive, red = true negative. The false positive rate for this classifier is 0.043. Accuracy for identifying sweeps vs non-sweeps is 0.958 and 0.848 for correctly identifying the specific class.

|  | | **Predicted Class** | | | | |
| --- | --- | --- | --- | --- | --- | --- |
|  |  | Hard | Linked-hard | Neutral | Linked-soft | Soft |
| **Simulated Class** | Hard | 1881 | 26 | 0 | 0 | 93 |
|  | Linked-hard | 65 | 1877 | 0 | 57 | 1 |
|  | Neutral | 0 | 0 | 1676 | 217 | 107 |
|  | Linked-soft | 6 | 294 | 220 | 1403 | 77 |
|  | Soft | 220 | 4 | 100 | 33 | 1643 |

Table S3. Simulated vs predicted sweep type classes for *P. t. ellioti* classifier on a second, independent set of simulations with identical parameters. We generated 2 x 10^3^ simulations for each class. Shaded cells reflect accuracy as a binary classifier: positive if hard or soft sweep, negative if linked sweep or neutral. Green = true positive, blue = false negative, purple = false positive, red = true negative. The false positive rate for this classifier is 0.042. Accuracy for identifying sweeps vs non-sweeps is 0.941 and 0.816 for correctly identifying the specific class.

|  | | **Predicted Class** | | | | |
| --- | --- | --- | --- | --- | --- | --- |
|  |  | Hard | Linked-hard | Neutral | Linked-soft | Soft |
| **Simulated Class** | Hard | 1847 | 69 | 0 | 3 | 81 |
|  | Linked-hard | 58 | 1911 | 0 | 30 | 1 |
|  | Neutral | 0 | 0 | 1611 | 278 | 111 |
|  | Linked-soft | 6 | 279 | 395 | 1243 | 77 |
|  | Soft | 186 | 2 | 196 | 69 | 1547 |

Table S4. Simulated vs predicted sweep type classes for *P. t. schweinfurthii* classifier on a second, independent set of simulations with identical parameters. We generated 2 x 10^3^ simulations for each class. Shaded cells reflect accuracy as a binary classifier: positive if hard or soft sweep, negative if linked sweep or neutral. Green = true positive, blue = false negative, purple = false positive, red = true negative. The false positive rate for this classifier is 0.031. Accuracy for identifying sweeps vs non-sweeps is 0.966 and 0.862 for correctly identifying the specific class.

|  | | **Predicted Class** | | | | |
| --- | --- | --- | --- | --- | --- | --- |
|  |  | Hard | Linked-hard | Neutral | Linked-soft | Soft |
| **Simulated Class** | Hard | 1911 | 43 | 0 | 0 | 46 |
|  | Linked-hard | 15 | 1904 | 0 | 81 | 0 |
|  | Neutral | 0 | 0 | 1360 | 522 | 118 |
|  | Linked-soft | 1 | 136 | 114 | 1700 | 49 |
|  | Soft | 140 | 1 | 68 | 43 | 1748 |

Table S5. Simulated vs predicted sweep type classes for *P. t. troglodytes* classifier on a second, independent set of simulations with identical parameters. We generated 2 x 10^3^ simulations for each class. Shaded cells reflect accuracy as a binary classifier: positive if hard or soft sweep, negative if linked sweep or neutral. Green = true positive, blue = false negative, purple = false positive, red = true negative. The false positive rate for this classifier is 0.014. Accuracy for identifying sweeps vs non-sweeps is 0.983 and 0.921 for correctly identifying the specific class.

|  | | **Predicted Class** | | | | |
| --- | --- | --- | --- | --- | --- | --- |
|  |  | Hard | Linked-hard | Neutral | Linked-soft | Soft |
| **Simulated Class** | Hard | 1950 | 13 | 0 | 0 | 37 |
|  | Linked-hard | 20 | 1949 | 0 | 31 | 0 |
|  | Neutral | 0 | 0 | 1769 | 181 | 50 |
|  | Linked-soft | 0 | 155 | 100 | 1731 | 14 |
|  | Soft | 123 | 1 | 49 | 19 | 1808 |
